# Shared long-term and short-term memory representational formats in occipital and parietal cortex

**DOI:** 10.1101/2021.02.16.431528

**Authors:** Vy A. Vo, David W. Sutterer, Joshua J. Foster, Thomas C. Sprague, Edward Awh, John T. Serences

## Abstract

Current theories propose that the short-term retention of information in working memory (WM) and the recall of information from long-term memory (LTM) are supported by overlapping neural mechanisms in occipital and parietal cortex. Both are thought to rely on reinstating patterns of sensory activity evoked by the perception of the remembered item. However, the extent of the shared representations between WM and LTM are unclear, and it is unknown how WM and LTM representations may differ across cortical regions. We designed a spatial memory task that allowed us to directly compare the representations of remembered spatial information in WM and LTM. Critically, we carefully matched the precision of behavioral responses in these tasks. We used fMRI and multivariate pattern analyses to examine representations in (1) retinotopic cortex and (2) lateral parietal cortex (LPC) regions previously implicated in LTM. We show that visual memories were represented in a sensory-like code in both tasks across retinotopic regions in occipital and parietal cortex. LPC regions also encoded remembered locations in both WM and LTM, but in a format that differed from the sensory-evoked activity. These results suggest a striking correspondence in the format of WM and LTM representations across occipital and parietal cortex. On the other hand, we show that activity patterns in nearly all parietal regions, but not occipital regions, contained information that could discriminate between WM trials and LTM trials. Our data provide new evidence for theories of memory systems and the representation of mnemonic content.

## Introduction

It is well-established that performance on a working memory (WM) or a long-term memory (LTM) task relies on the partial reactivation, or reinstatement, of patterns of cortical activity that were present during the initial perception of the remembered item (D’Esposito & Postle, 2015; Serences, 2016; Squire & Wixted, 2011; Xue, 2018). For example, viewing a blue couch elicits feature-specific patterns of activity in visual regions selective for color and complex objects. Then, when trying to maintain the colored object in WM, or retrieve it from LTM, patterns of neural activity that resemble those at encoding are reinstated. That said, evidence for feature-specific cortical reinstatement has typically been assessed separately in studies of LTM (Bosch, Jehee, Fernández, & Doeller, 2014; Favila, Samide, Sweigart, & Kuhl, 2018; Kuhl & Chun, 2014; Ritchey, Wing, LaBar, & Cabeza, 2013; Xiao et al., 2017) and studies of WM (Emrich, Riggall, Larocque, & Postle, 2013; Ester, Anderson, Serences, & Awh, 2013; Ester, Sutterer, Serences, & Awh, 2016; Harrison & Tong, 2009; S.-H. Lee, Kravitz, & Baker, 2013; Riggall & Postle, 2012; Serences, Ester, Vogel, Awh, & Serences, 2009; Sprague, Ester, & Serences, 2014, 2016). While several prominent theories suggest that both WM and LTM retrieval rely on shared neural substrates (Atkinson & Shriffin, 1968; Cowan, 1995; Kosslyn, 1980), we are unaware of any direct tests of this hypothesis, perhaps because it is challenging to match the WM and LTM tasks on important dimensions such as overall task performance.

Here, we strived for a direct comparison of cortical reinstatement during WM and LTM. Participants either maintained a spatial position in WM or retrieved it from LTM and maintained it for the same duration while we measured responses with functional magnetic resonance imaging (fMRI). Notably, the remembered information was the same across conditions – it simply arose from a different source (perception for WM vs. an internal representation for LTM). We used multivariate pattern analysis to examine the remembered representations in early occipital visual areas and across several regions in retinotopic and lateral parietal cortex. The 8 retinotopic regions we defined (V1 through dorsal parietal regions IPS2) are known to have sensory-like WM representations (Emrich et al., 2013; Ester et al., 2016; Riggall & Postle, 2012; Sprague et al., 2014, 2016) that are finely resolved enough to decode stimulus features such as orientation or motion direction. That is, a model trained on sensory representations can decode feature information during a WM task. Recent evidence suggests that in IPS, feature-specific WM representations can be encoded in a non-sensory format (Rademaker, Chunharas, & Serences, 2019). That is, only a model trained on WM representations can decode information during the memory task. As for LTM, the data on reinstated stimulus features in retinotopic regions is limited to V1-V3 (Bosch et al., 2014) or relies a single large occipitotemporal ROI (Favila et al., 2018). Therefore we also examined a set of lateral parietal regions (dorsal and ventral lateral IPS; angular gyrus), which have been shown to contain feature-specific information during retrieval from LTM (Kuhl & Chun, 2014; H. Lee, Chun, & Kuhl, 2016). Their role in WM is not well understood (Favila, Lee, & Kuhl, 2020; Favila et al., 2018; Xiao et al., 2017). Lastly, we took advantage of our experimental design to test whether the reinstated information in these regions is identical between WM and LTM or if the feature-specific information is maintained via different representational format.

An overview of the WM and LTM tasks is shown in **Figure 1**. Importantly, we carefully matched behavioral performance across memory tasks and minimized any differences in visual properties. We then used an inverted encoding model (IEM) to quantify the fidelity of cortical reinstatement in both tasks, based on the amount of spatial information in fMRI activation patterns. To test whether the remembered spatial position was represented in a sensory-like format, we trained the IEM on an independent perceptual task and tested it on either the WM maintenance task or LTM retrieval task. To test if a mnemonic representation was in a non-sensory-like format, we trained the IEM on one of the memory tasks instead. First, we found that memories for spatial position maintained in WM or retrieved from LTM were represented in a sensory-like format across all retinotopic regions of interest (ROIs). Furthermore, the fidelity or quality of these representations was statistically indistinguishable between WM and LTM. We found that lateral IPS and angular gyrus both tracked some spatial information held in memory. However, this information was represented in a non-sensory format.

**Figure 1.**
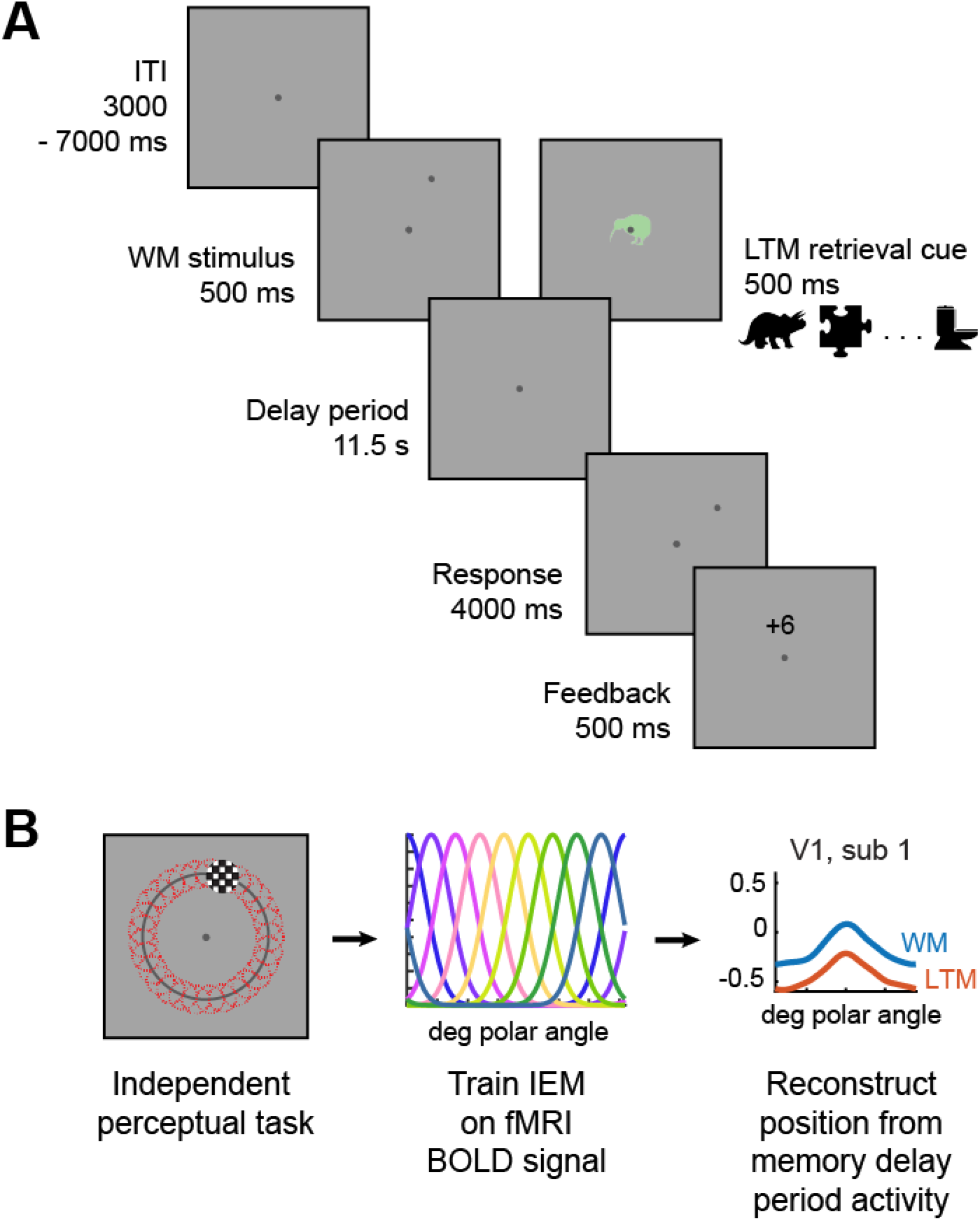
Memory tasks and perceptual task presented to participants in the scanner. (A) The two memory tasks were identical except for the memory cue – in the WM task, participants saw a dot indicating the position along an annulus, along with an irrelevant clip art item; in the LTM task, participants instead saw a previously studied clip art item that they had learned to associate with a spatial position during behavioral training. After an 11.5s delay period, participants reported the remembered spatial position by rotating a randomly placed dot to the correct position. (B) An independent task was used to map the position selectivity of voxels in retinotopically defined regions of visual and parietal cortex. Participants detected an occasional dimming of a checkerboard stimulus that appeared at randomly ordered locations along a ring. This independent task was used to train the IEM, which was then tested on the memory tasks.

Next, we used a 2-way linear classifier to decode the memory task that subjects were performing on each trial. We found that the memory task could be decoded from activation patterns in parietal regions (both retinotopic and lateral) but not in early visual cortex. Our results provide direct evidence that WM and LTM rely on similar neural representations in occipital and parietal cortex. However, parietal cortex, unlike early visual cortex, also encodes information about the source of the remembered information in addition to coding the location of remembered positions.

### Methods

#### Participants

We trained 12 human participants (8 female) on both the LTM and the WM task. Two participants dropped out of the study during behavioral training. This left 10 participants (6 female) from which we collected both behavioral and fMRI data, 2 of which were co-authors on this paper. Six of these participants (participant IDs BI, BJ, BO, and BC) had previously completed a set of retinotopic mapping scans in the lab for other studies (Henderson & Serences, 2019; Rademaker et al., 2019). For the remaining 4 participants, we collected new retinotopy data to define each region of interest (see *Region of interest definition*). All participants provided written informed consent. Participants were compensated for their time ($20/hour for fMRI, $10/hour for behavior) unless they were paper co-authors. These procedures were approved by the local UC San Diego Institutional Review Board.

#### Pre-scan behavioral training

We trained participants to form long-term memory pairings between 24 unique clip art items and 24 unique spatial positions (**Figure 1**). After one day of LTM study and retrieval, we continued LTM training while also measuring their performance on an analogous working memory task. In both tasks, participants were asked to report the remembered spatial positions as precisely as possible. To minimize differences in effort and difficulty of recall between the two memory tasks at the time of scanning, participants were trained until their error on the LTM task was stable.

For each participant, the 24 spatial positions were selected randomly from 24 bins evenly spaced along an isoeccentric ring 3.9° visual angle from a central fixation point. As a result, the set of locations was roughly uniformly distributed around a circle for each participant, and each participant had a slightly different set of positions. Each spatial position was rendered as a dot subtending 0.2° visual angle (in polar coordinates of the 360° circle, this subtended 1.4° polar angle). The clip art items (diameter 1.5° visual angle) were filled outlines of real-world objects taken from a bank of 48 images that all subtended the same area of retinal space (**Figure 1**). These stimuli have been used in previous LTM studies, and are all royalty-free images (Sutterer & Awh, 2015; Sutterer, Foster, Serences, Vogel, & Awh, 2019).

The LTM study-retrieval task on the first day of training consisted of 6 alternating runs of a study phase and a retrieval phase. During the study phase, the participant was shown each clip art item at the center of the screen and its associated spatial position. The trial order was randomized. This study phase was self-paced – while each pairing was always presented for 500 milliseconds (ms) followed by at least a 1000 ms blank period, participants chose when to advance to the next item by hitting the space bar. During the retrieval phase of the task, each clip art item was presented at the center in a randomized order, and participants were asked to report the associated spatial position as precisely as possible by clicking on a part of the visible isoeccentric ring. After each retrieval trial, participants received two forms of feedback about their performance: the true position was shown in dark gray on the ring, and the error was shown as a signed integer at the center. This number indicated how far their report deviated from the true position (+ if report was too far clockwise, - for counter-clockwise).

On all remaining days of pre-scan training, participants practiced both the LTM retrieval task and a similar WM maintenance task (similar to **Figure 1A**, with different timing). The order of the tasks was counterbalanced across participants. In the LTM retrieval task, the participant was shown one of the 24 clip art items at the center (500 ms). After a delay (1000 ms), they were presented with the response probe: a dot appeared at a random position along the (now invisible) isoeccentric ring, and participants used four keyboard keys to rotate this dot along the ring until it matched the item’s associated position as precisely as possible. The two outer buttons moved the dot along the ring at a fast rate, whereas the two inner buttons moved it along a slower rate. This allowed participants to make fast but precise responses. After the 4000 ms response period, participants received numerical feedback on their response error (**Figure 1**). Unlike the first day of training, this was the only form of feedback. The intertrial interval varied between 200 and 500 ms. Participants repeated this set of 24 trials per run, for a total of 6 runs in the LTM task block.

The WM maintenance task was nearly identical to the LTM task in timing and presentation. The one exception was that the initial memory array displayed a dot at a random spatial position on the ring, in addition to a clip art item presented at the center that was randomly selected from a separate set of 24 unique clip art items. Participants were asked to remember the position of the dot during the 1 second delay, and to use the keyboard keys to rotate the response dot to the position of the memorandum (**Figure 1**). The clip art item was a task-irrelevant sensory control, and there was no predictive relationship between WM position and clip art item. As with the LTM task, a set of 24 trials was repeated 6 times.

All participants performed 6 runs of each task on each training day. The first two participants (BG, BH) completed more training runs as we piloted the best way to collect response data (mouse or keyboard). Their behavioral performance in the scanner did not appear significantly different from the other subjects. All participants were trained on both the LTM and WM tasks until their mean recall error on the LTM retrieval task was stable and no longer decreasing. One participant (BV) requested to repeat the initial LTM study training on day 4 of training.

#### Behavioral tasks performed in fMRI scanner

Participants performed both LTM and WM tasks in the scanner (**Figure 1A**). While these were very similar to the behavioral training task, they differed in several key respects: (1) instead of a 1000 ms delay between the cue and the response, we inserted a 11,500 ms delay to accommodate the slow hemodynamic response; (2) LTM and WM runs were alternated instead of blocked, in the order preserved from the training; (3) participants viewed the experiment through a head-coil mounted mirror pointing at a gray rectangular screen (120×90 cm) at the foot of the scanner bore (∼3.85 m viewing distance); (4) responses were made via a 4-key button box; and (5) the intertrial intervals were longer (uniformly distributed in 200 ms steps between 3000 and 7000 ms), to aid in estimating the stimulus-evoked BOLD response. Participants continued to receive feedback on every trial that indicated the signed response error.

In addition to the memory tasks, participants also performed an independent mapping task which we used to train the spatial encoding model (see below) and to localize voxels responsive to the isoeccentric ring (as before, 3.9° from central fixation point). In this task, participants fixated at a central point as a flickering stimulus (6 Hz) appeared at some location on the screen for 3000 ms. Participants monitored the flickering stimulus for an occasional contrast change (8 of 54 total trials per run), and reported whether the contrast increased or decreased on those trials (pedestal at 70% Michelson contrast). On most trials, the stimulus was a circular checkerboard (0.9° radius, 1.36 cycles/deg) centered at a point on the isoeccentric ring. Each location was pseudorandomly drawn from one of 30 evenly spaced polar bins. On 5 trials, participants instead saw a ring aperture (1° wide) with a similar flickering checkerboard pattern (3.9° from center). On the target trials, either the entire circular checkerboard changed contrast, or a 0.681° (36° polar angle) wide area of the ring changed contrast for 500 ms. We also included 6 null trials to aid in the estimation of the stimulus-evoked BOLD response. Trial order was randomized, with inter-trial intervals (ITIs) varying between 2000 and 5000 ms (mean 3,500 ms). The magnitude of the contrast change was manually adjusted by the experimenter on each run to keep task performance for each participant near 75% (participant averaged mean and bootstrapped 95% CIs: 71.7% accuracy [60.11%, 81.62%]; Michelson contrast change 0.50 [0.42, 0.58]).

#### Statistical analysis

The majority of the statistical analyses reported in the paper and all confidence intervals (CIs) were generated by bootstrapping. We opted for a non-parametric approach because it carries fewer assumptions about the distribution of the underlying data.

Unless otherwise noted, the bootstrapping was done by resampling with replacement across the N = 10 participants for 10000 iterations. To correct the confidence intervals for a bias towards narrowness in small samples, we drew N-1 samples on each iteration (Hesterberg, 2011). Then the mean of the value of interest (e.g., response error) was computed on each resampling iteration to generate a distribution of means for the sampled population. Generally, we calculated 95% CIs using the percentile method. When computing the fidelity metric, however, we also corrected for skewness in the CIs across participants by applying the bias correction and acceleration (BCa) adjustment (Davison & Hinkley, 1997; Hesterberg, 2014) after estimating bias and acceleration by jackknife. BCa-adjusted values also produce an accompanying p-value.

To evaluate the evidence for the null hypothesis that there was no difference between the WM and LTM tasks, we computed a Bayes factor for the comparison between conditions. We used the *BayesFactor* package in R 3.5.3 to convert a paired-sample t-value into a Bayes factor, given a standard Cauchy prior on the effect size and Jeffreys prior on variance (Rouder, Speckman, Sun, Morey, & Iverson, 2009). For a stable estimate of the t-value in our small sample, we computed a t-value on every iteration of the across-subject bootstrap described above. This generated a distribution of t-values. We took the mean t-value of the bootstrapped distribution to calculate the Bayes factor in favor of the null hypothesis of no difference between WM and LTM (BF_01_).

To assess effects across ROIs, we ran repeated-measures ANOVAs on either the representational fidelity metric or the decoding accuracy. Since these measures are not guaranteed to be normally distributed, we compared the F-score from a typical ANOVA to a null distribution of F-scores that was generated by permuting the data across all factors (e.g. ROI, temporal epoch, and memory task), separately within each participant, for 10,000 iterations (Manly, 2007).

When multiple statistical comparisons are made (i.e. repeated for each ROI), we corrected across all test p-values using the false discovery rate procedure (q = 0.05 unless otherwise noted; Benjamini & Yekutieli, 2001).

#### Behavioral data analysis

To describe participant performance on each memory task, we examined each participant’s trial-by-trial response error. This is the signed difference between the true location of the memorandum and the participant’s response (between 0 and ±180°). Previous findings have shown that a histogram of response errors can be described as a mixture of a uniform distribution, representing guesses, and a circular Gaussian distribution centered near the correct response (Zhang & Luck, 2008). The two parameters of the circular Gaussian describe the mean response (µ), which describes any systematic bias, and the standard deviation of the responses (*SD*), which describe the variability of the memory reports. These parameters were fit using publicly available code (Schneegans & Bays, 2016).

All behavioral data analyses used 10,000 resampling iterations.

#### Magnetic resonance imaging

We obtained all structural and functional MR images from participants using a GE 3T MR750 scanner at University of California, San Diego. We collected all functional images (19.2 cm ^2^ FOV, 64 x 64 acquisition matrix, 35 interleaved slices, 3 mm ^3^ voxels with 0 mm slice gap, 263 volumes per memory run and 179 volumes per mapping run) using a gradient echo planar pulse sequence (2000 ms TR, 30 ms TE, 90° flip angle) and a 32-channel head coil (Nova Medical, Wilmington, MA). Five dummy scans preceded each functional run. A high-resolution structural image was acquired at the end of each session using a FSPGR T1-weighted pulse sequence (25.6 cm ^2^ FOV, 256 x 192 acquisition matrix, 8136/3172 ms TR/TE, 9° flip angle, 1 mm ^3^ voxels, 172 volumes). All functional scans were co-registered to the anatomical images acquired during the same session, and this anatomical was in turn co-registered to the anatomical acquired during the retinotopy scan.

EPI images were unwarped with a custom script from UCSD’s Center for Functional Magnetic Resonance Imaging that calls the FSL PRELUDE & FUGUE functions. All subsequent preprocessing was performed in BrainVoyager 2.6.1, including slice-time correction, affine motion correction, and temporal high-pass filtering to remove slow signal drifts over the course of each session. Data were then transformed into Talairach space and resampled to a 3×3×3 mm voxel size. Finally, the BOLD signal in each voxel was Z-transformed on a scan-by-scan basis. All subsequent analyses were performed in MATLAB using custom scripts.

For the encoding and decoding analyses, we balanced the data by equating the number of runs for each memory task. Some data was discarded due to excessive movement (>3 mm in one run/scan). In these cases, we dropped runs until there were an equal number of WM & LTM runs.

#### Region of interest definition

The retinotopic regions of interest (ROIs) were first defined for each participant following a previously published procedure (Sprague & Serences, 2013). Briefly, we collected fMRI data in a separate 1.5 hour session that mapped the visual field. While one task was a passive viewing task, the other task required participants to attend to the portion of the visual field that contained the checkerboard stimulus and report an occasional dimming in one portion of the stimulus. This attention task allowed us to reliably define parietal ROIs. In total, we defined 8 retinotopic regions: V1, V2, V3, V4, V3A/B, IPS0, IPS1, and IPS2.

We then applied a mask to the retinotopically-defined occipital and parietal regions to isolate voxels that show some response to the area where the spatial positions were presented. This functional localizer relied on defining different conditions for the circular checkerboard and ring aperture checkerboard trials during the mapping task. We estimated BOLD response activation to each condition by convolving the trial events with a canonical two-gamma HRF (peak at 5 s, undershoot peak at 15 s, response undershoot ratio 6, response dispersion 1, undershoot dispersion 1). We then solved a generalized linear model (GLM) on all runs of this mapping task for a single participant. This produced a statistical parametric map of voxels with a significant BOLD response change attributable to each condition (FDR q = 0.05, Benjamini and Yekutieli, 2001). The localizer mask only included voxels that responded with a significant BOLD response increase during the ring aperture condition.

In addition to retinotopic regions, we also defined three lateral parietal regions: dorsal lateral intraparietal sulcus (dLatIPS), ventral lateral intraparietal sulcus (vLatIPS), and angular gyrus (AnG). We defined these regions following a previous method used in a study of episodic memory (Favila et al., 2018). First, we used the Freesurfer parcellation to specify the lateral parietal cortex as any area in the superior parietal, inferior parietal, and supramarginal parcels. We then subdivided this portion of cortex using a 17-network atlas defined from resting state functional connectivity data (Yeo et al., 2011). dLatIPS was part of network 12 in the atlas, and vLatIPS was part of network 13. Both of these networks are part of the frontoparietal control network. The AnG was defined by combining the small parietal nodes from networks 15, 16, and 17, which are part of the default mode network.

Note that these atlas-defined regions were not defined using any functional data. As a result, they exhibit some overlap with the retinotopically defined parietal regions (see **Figure 2**).

**Figure 2.**
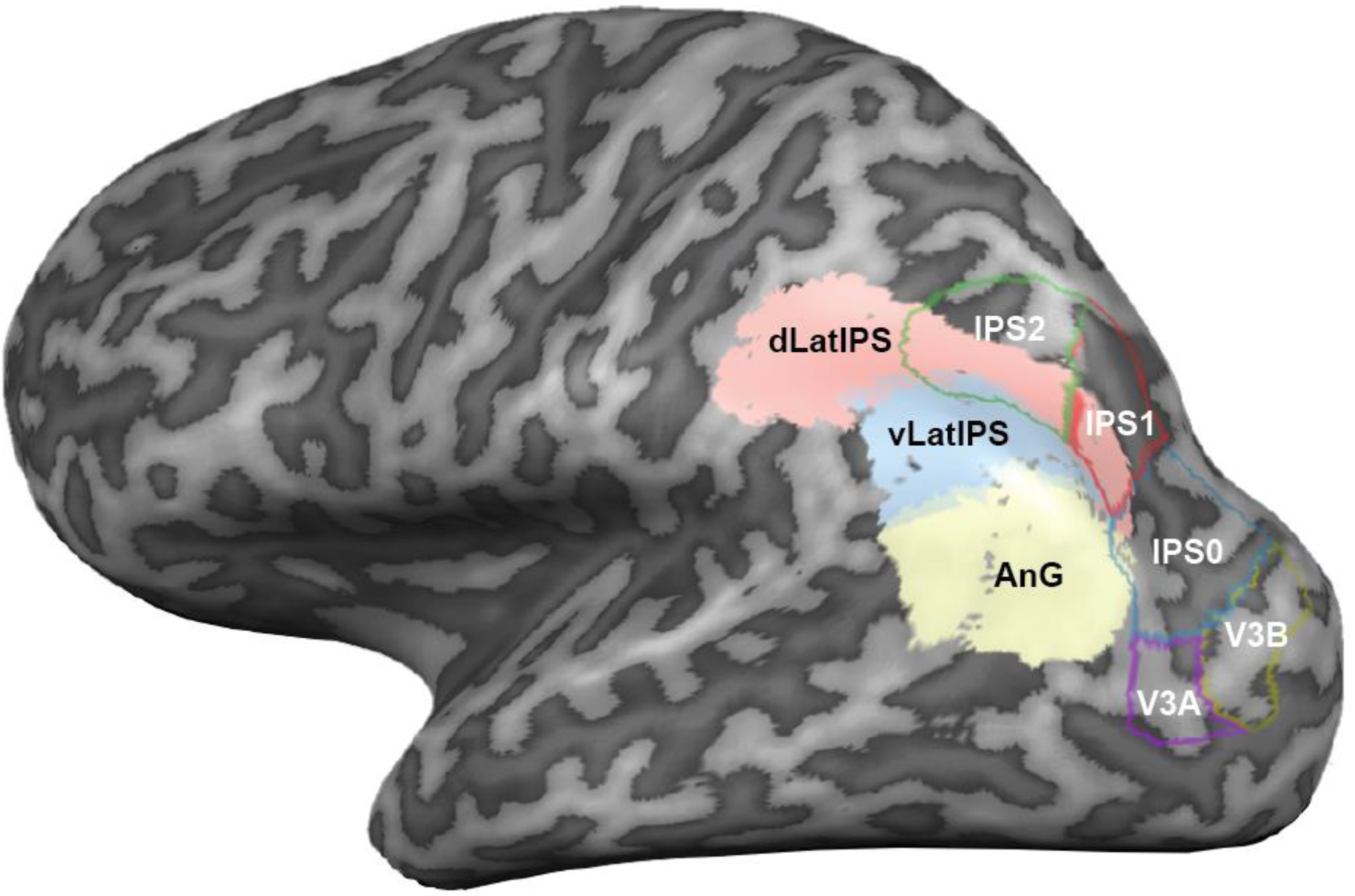
Retinotopic parietal ROIs (outlines, white labels) and lateral parietal ROIs (filled, black labels) for the left hemisphere of an example participant.

### Inverted encoding model for spatial position

Following previously published methods, we first estimated an encoding model for spatial position using z-scored BOLD responses obtained from the independent perceptual task (Brouwer & Heeger, 2009; Sprague et al., 2018; Sprague & Serences, 2013). This allowed us to estimate the response of each voxel to any arbitrary spatial position covered by the visual stimuli.

We defined 9 evenly spaced spatial channels, or basis functions, along the polar angle axis (**Figure 1**). This was the only axis we varied in our mapping runs. Each spatial channel was defined by Equation 1, assuming a discretized polar axis (i.e. 0 to 2π in steps of 0.0175 radians, or 1 degree). The width and center of the basis functions are modified by the term 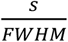, where *s* is the circular distance between the channel center and the polar axis, and FWHM is the desired full-width half-maximum of the response of the spatial channels. Our basis functions had a FWHM of 60°. Finally, to keep the encoding model weights within an interpretable range, we want each spatial channel to have a baseline of 0 and a maximum of 1. To achieve this, we used a positive, half-wave rectified cosine.

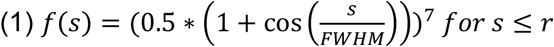

Finally, although cosine functions are periodic, each spatial channel should only have a peak response at its center, and nowhere else. We mask the function by setting all values beyond the radius *r* to 0 (here, r = 151).

The basis set **S** is the 9 spatial channels in the discretized polar axis (*c* channels x *p* points along the polar axis, here 9 x 360). For every training trial, we used the basis set to specify a modeled activation on each of the *c* channels for each presented mapping stimulus, resulting in a design matrix **C** (*c* channels x *n* trials).

The forward encoding model is then described in Equation 2, where B = BOLD responses (*v* voxels x *n* trials). As long as *n* > *c*, it is possible to solve for **W** using the pseudoinverse (Equation 3). We solved Equation 3 using *mldivide* in MATLAB.

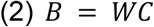

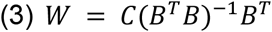

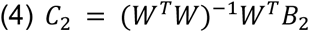

After training the model by solving for **W**, we inverted the encoding model (Equation 3) to estimate the activation of each spatial channel, given the BOLD responses on a separate test dataset and the independently trained model from the same participant (Equation 4). The spatial channel activations allowed us to estimate the spatial position of the stimulus that was presented in each trial. The test dataset could be the BOLD responses of voxels during the memory delay of either the WM or LTM task (Ester, Sprague, & Serences, 2015; Sprague et al., 2014), or the BOLD responses to a held out portion of the independent mapping task. Notably, we attempted to recover spatially-specific information in the patterns of voxel responses during each of two memory tasks when *no visual stimulus was shown on the screen*. Furthermore, we used a ‘fixed’ encoding model estimated using a separate dataset and task, enabling us to directly compare model-based representations across memory tasks (Sprague et al., 2018; Sprague, Boynton, & Serences, 2019).

We estimated channel responses **C_2_** from two different types of BOLD data **B_2_**. In **Figure 4, B_2_** consisted of the BOLD response at each TR (after motion-correction and z-scoring within each run), for a total of 8 TRs. In other analyses described below, we averaged the data across TRs for use in the inverted encoding model.

To generate a continuous representation of spatial position, we multiplied **C_2_** by the basis set **S**. We re-centered all of the model-based representations such that the memory stimulus is at 0°, and average across all trials. These are the curves plotted in **Figure 4A**.

For **Figure 5**, we varied the dataset used for training and testing the inverted encoding model. To identify which regions contained significant representations of the sensory stimuli (i.e., the spatial position), we trained and tested within the mapping dataset with leave-one-run-out cross-validation. To characterize whether a region contained significant spatial representations within a memory task, we trained and tested within the memory task, only using TRs from the late delay period. To ensure that the datasets are balanced across spatial position, we randomly resampled from each of 24 position bins and held out those 24 trials for testing. This resampling procedure was repeated 1000 times, and the results were averaged across these iterations.

Finally, to test whether a region encoded spatial representations similarly across memory tasks, we trained on one memory task and tested on the other, again only using TRs from the late delay period.

#### Representational fidelity metric

To quantify the model-based memory representations, we compute a metric which describes the fidelity of the spatial representation. This is similar to previously published metrics but is modified so that it monotonically increases as the representation decreases in width (Sprague et al., 2016; Wolff, Jochim, Akyurek, & Stokes, 2017). Consider each point of the spatial IEM-based representation in polar coordinates as [*r_n_, θ_n_*], where *r* is the value of the representation in arbitrary units (y-axis in **Figure 4A**) and *θ* is the polar angle (x-axis in **Figure 4A**). The representational fidelity is directly proportional to the average directional energy of the spatial representation pointing in the expected direction (0°). We computed this by finding the mean vector for across all *n* points using two metrics of circular data, the average preferred direction and the dispersion (Jammalamadaka & SenGupta, 2001). Since these metrics only exhibit the desired behavior when *r* is positive, and because we were focused on fidelity as opposed to baseline offsets, we forced the minimum of the model-based representation to be 0 by adding an offset to every representation.

First we found the mean direction of the spatial IEM-based representation (i.e. the angle of the vector) using the *circ_mean* function in the CircStats MATLAB toolbox (Berens, 2009). We then computed how closely the vector points to the expected value of 0 by taking the cosine of this value.

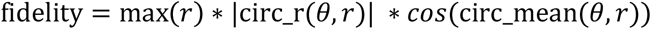

We then calculated the dispersion of the spatial representation (i.e., the vector length), which scales monotonically with the width of the function. As the spatial representation becomes narrower and more precise, the vector length increases. The dispersion is computed with *circ_r* (Berens, 2009).

Finally, we multiplied the mean direction by the dispersion and scaled the result by the maximum value of the spatial representation, yielding a fidelity metric which monotonically increases as a given spatial representation increases in amplitude and in precision (**Figure 4B**). The metric is negative when the mean vector points in the opposite direction (180°), and close to 0 if the spatial representation is flat and contains no information about the remembered location.

As noted in the *Statistical analysis* section, we resampled across participants to generate confidence intervals for the sampled population. To generate single-participant confidence intervals, we bootstrapped the data by resampling across trials. If the 95% CIs overlapped with 0, then there was no significant memory representation for that participant or for that region. When we compared fidelity across memory tasks, we subtracted the two bootstrapped distributions (e.g. WM – LTM) and compared this difference distribution against 0.

#### Support vector machine classification

The analyses described above allowed us to generate continuous estimates of remembered stimuli to precisely compare the quality of spatial memory representations across ROIs and memory tasks. We were also interested in whether activation patterns in these ROIs could discriminate between the two memory tasks. To test this, we trained a support vector machine (SVM) with the default linear kernel to discriminate between WM trials and LTM trials. This was done using MATLAB’s *fitclinear* routine.

We first divided the data into two temporal epochs during the memory delay period and averaged across the TRs in those epochs (early epoch: TRs 2 – 4, late epoch: TRs 5 – 7). Classifier training and testing were performed separately for each epoch, ROI, and participant. To ensure that classification performance was not driven by a global difference in univariate response between the memory tasks, we computed the global mean across all trials and voxels for each memory task. We then subtracted this constant from the data before we performed any of the SVM analyses below.

Mean classifier accuracy was determined by k-fold cross-validation (k = 12, which is the typical number of runs in each task condition) using MATLAB’s *crossval* function. We then assessed statistical significance by randomizing the labels associated with each task and generated a null distribution of classifier accuracies over 1000 iterations.

To combine the data across participants, we took all classifier accuracies for a given ROI and temporal epoch (10 values, one for each participant) and calculated the mean. We then performed a similar operation on the permuted null distributions, concatenating all the distributions (10 participants x 1000 iterations) and calculating the mean across participants to generate a single null distribution for the mean.

We computed empirical one-tailed p-values by comparing when the mean classifier performance was higher than the permuted null values (1 – mean(classAcc > nullDistr)). Note that we used one-tailed tests to reflect the fact that the classifier accuracies were not predicted to drop significantly below chance. These p-values were subject to FDR-correction across temporal epochs and ROIs. We used a stricter FDR q = 0.025 to reflect the fact that we only used one-tailed p-values.

## Results

### Behavioral data

Outside of the scanner, participants were trained on both the LTM and WM tasks until their mean recall error on the LTM retrieval task was stable and no longer decreasing (mean number [95% CIs] of training days 7.39 [5.78, 9.22]). However, this training target did not enforce similar performance between tasks (participant averaged error on the last day: WM 4.29° [3.58°, 5.12°]; LTM 6.54° [4.69 °, 8.52°]). To characterize each participant’s recall success and precision, we plotted a histogram of their recall errors and fit the data with an additive mixture of a circular Gaussian and a uniform distribution (Harlow & Yonelinas, 2016; Zhang & Luck, 2008). The standard deviation (*SD*) of the Gaussian is a measure of the spatial precision of their recall, with larger SDs corresponding to imprecise recall. The height of the uniform distribution is a measure of their rate of forgetting. Here we report the complement of this number, the probability of recall (*P*(recall)). The interval training procedure combined with precise feedback significantly increased both the precision and probability of recall in the LTM task (difference between first and last set of trials: *SD* 14.12° [2.88°,37.23°], p < .001; *P*(recall) 0.05 [0.01,0.10], p = 0.01). Although our training procedure did not require that performance was matched between the WM and LTM tasks, we nevertheless found that mean response error was comparable between the tasks during the sessions in the scanner (participant averaged WM error 7.40 [5.67, 9.53]; LTM error 7.78 [4.54, 11.88], p = 0.78; Bayes factor in favor of null BF_01_ = 3.24). The probability of recall was high (*P*(recall): WM 0.98 [0.96, 1.00], LTM 0.97 [0.94, 1.00]; **Figure 3b, right**), and similar between both memory tasks (p = 0.35; BF_01_ = 2.40). As a group, participants also had similar recall precision between both tasks (mixture model *SD*: WM 7.61 [6.56, 8.69], LTM 7.91 [5.38, 10.81]; p = 0.85; BF_01_ = 3.23). However, recall precision varied more on the LTM task: some participants had worse precision in the LTM task than the WM task, while others had better precision (**Figure 3b, left**).

**Figure 3.**
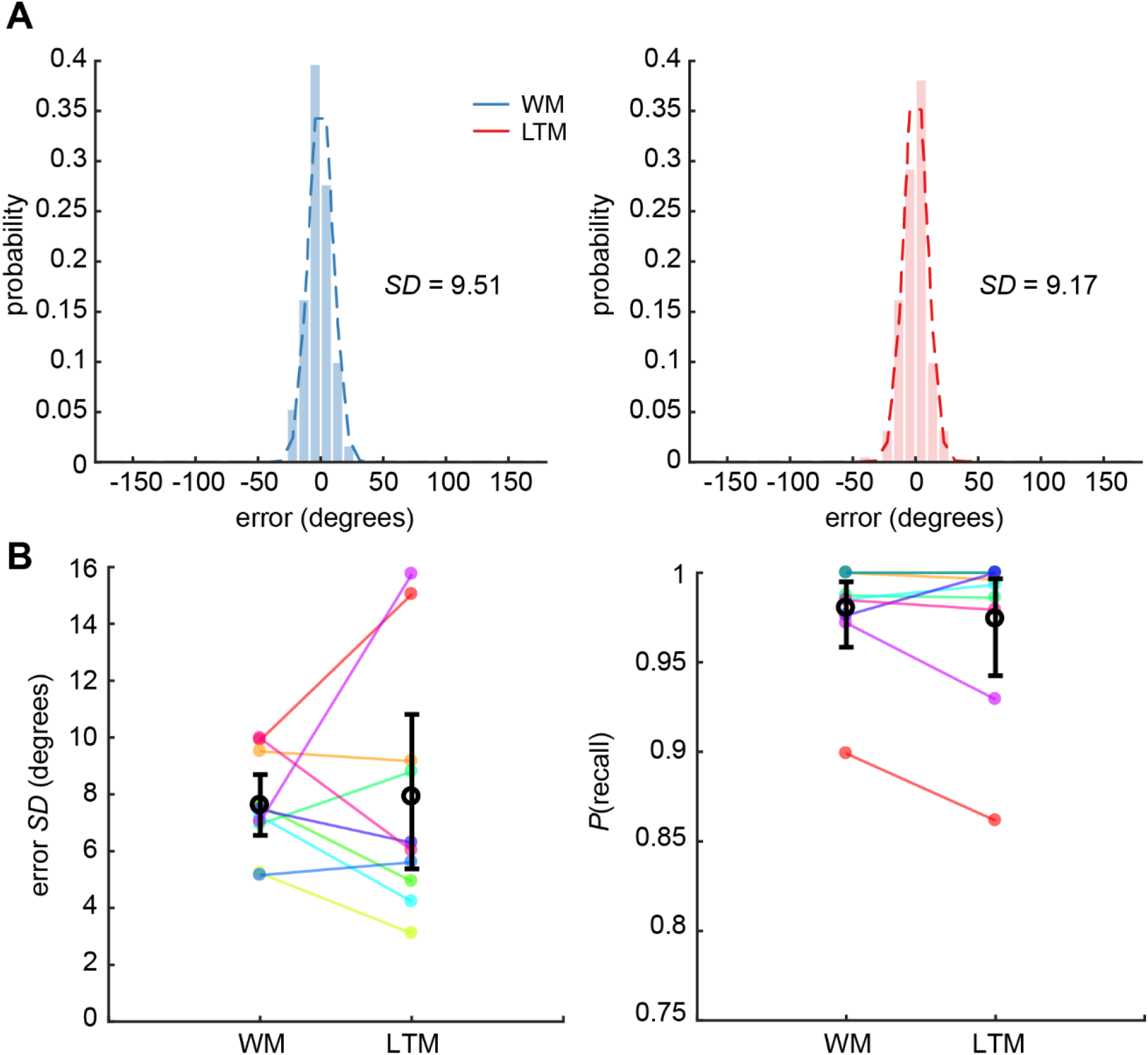
Memory recall across both scanner tasks is similar. (A) The distribution of memory recall errors for an example participant. We fit a mixture model to this distribution for every participant, where the Gaussian distribution characterized the variability of recall (standard deviation, SD) and the uniform distribution characterized the likelihood of recall (P(recall)). (B) The mixture model fit parameters for each subject and task. The mean across participants and 95% CIs are shown in black.

**Figure 4.**
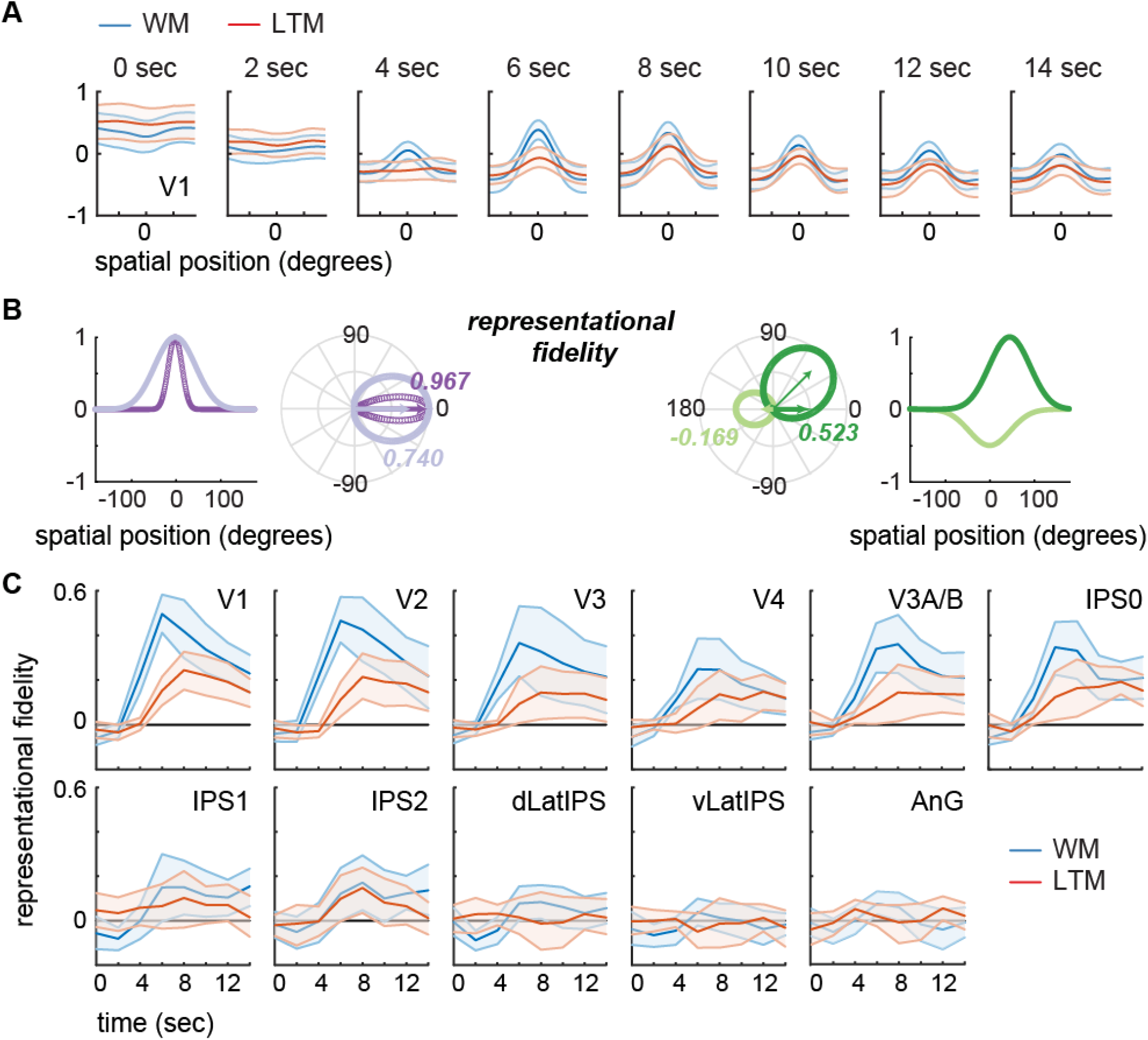
Model-based representations of remembered spatial positions over time for both memory tasks. Error bars are participant-resampled 95% CIs. (A) The timecourse of model-based spatial representations using V1 data, averaged across participants (0 sec is stimulus onset). Remembered position was represented similarly for WM and LTM late in the delay period (8 – 14 seconds). (B, left panel) Example spatial representations and their corresponding fidelity values, a single number which characterizes the quality of tuning toward the remembered position (always set to 0° here). Each spatial representation was plotted in polar space. The fidelity metric is equivalent to the length of the bold horizontal vectors. Dark purple: the best model-based spatial representation is narrow and centered exactly at 0°. Light purple: a broader spatial representation has a shorter mean vector, capturing the fact that less ‘energy’ is at 0°. Dark green: a spatial representation that is slightly offset from zero has a short x-component compared to a represented centered at zero, and therefore a lower fidelity value. Light green: an inverted spatial representation has a mean vector that points in the opposite direction, resulting in a negative fidelity value. (B, right panel) Representational fidelity timecourses for all retinotopic areas we analyzed. CIs that intersect with 0 are not significant. dLatIPS: dorsolateral IPS; vLatIPS: ventrolateral IPS; AnG: angular gyrus.

**Figure 5.**
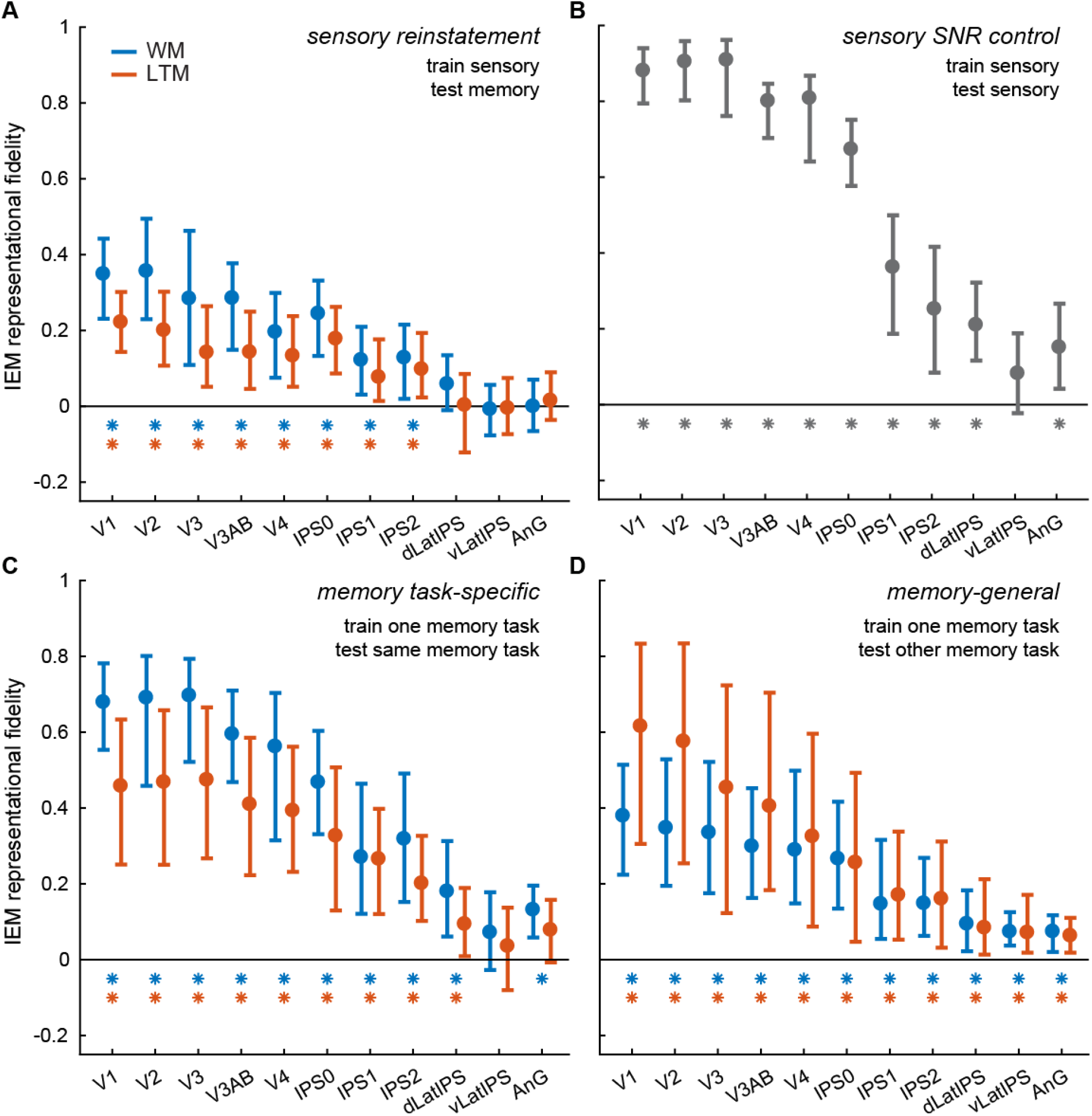
Model-based representations for different combinations of training and testing data, taken from the end of the delay period (8-12 seconds). For each plot, we ran a permuted 2-way ANOVA of ROI by memory task (or for the sensory control, a 1-way ANOVA across ROIs). In all 4 cases, there was a significant main effect of ROI (p < .001). We found a significant 2-way interaction for both the independent training set (p = .003) and the cross-training procedure (p = .001), but no main effect of task. Training within memory tasks yielded a borderline significant effect of memory task (p = .052), as the WM task generally resulted in higher fidelity than the LTM task. Asterisks indicate significant information after FDR-correction across all ROIs and both memory tasks, q = 0.05.

### Sensory-like representations of remembered spatial position

To assess how spatial information was encoded during the delay period of both memory tasks, we trained an encoding model using data from an independent perceptual task, which allowed us to map the spatial selectivity of visually responsive voxels in several retinotopic regions of interest (ROIs). Having trained the model of the independent perceptual task, we inverted the encoding model to reconstruct spatial representations of the remembered location from the pattern of activity across voxels in the WM or LTM tasks. If this reconstruction is highly accurate, then we can conclude that the pattern of voxel activations during the perceptual task is similar to the pattern of activations during the delay period of the memory tasks (i.e. it is sensory-like). This is a strong test of the cortical reinstatement hypothesis, which posits that the neural representation of an item during perception is reinstated when that item is recalled from memory. Furthermore, using an encoding model trained on a single independent task allows us to directly compare between spatial representations across WM and LTM (Sprague et al., 2018, 2019).

The spatial reconstructions show activation at each modeled location, and peaks within reconstructions can be interpreted as visual spatial memory representations that can be parameterized as a curve in circular (polar) coordinate space since the eccentricity of the stimuli was fixed (**Figure 4B**). To average across trials with different remembered positions, we re-centered the reconstruction on each trial so that the remembered position was at 0 degrees. In **Figure 4A**, we plot these averaged spatial reconstructions from area V1 at several timepoints during the memory delay of both tasks. As expected, spatial information during the delay period did not emerge until 4-6 seconds after stimulus onset, when the BOLD response evoked by a stimulus is expected to peak. Additionally, the spatial reconstruction was higher amplitude in the WM task than in the LTM task early in the delay period. This early difference is likely due to fact that the remembered position was marked by a dot in the WM task, but not the LTM task (Figure 1a), which should evoke a sensory response. However, late in the delay period (∼10 seconds after stimulus onset), when lingering sensory activation should have decayed, we found that model-based spatial representations were similar between the WM and LTM tasks.

To quantify the model-based spatial representations, we computed a metric to measure the representational fidelity of each re-centered spatial representation (*Methods*). We illustrate some properties of this fidelity metric and its computation in **Figure 4B** (left panel). First, each point in the spatial representation is replotted in polar space. We then find the mean vector of all of these points, which describes the average directional energy of the spatial representation. The fidelity, then, is proportional to the length of the vector along the expected direction (i.e., 0°, or the x-component of the vector; bold italic numbers in **Figure 4B**, left panel). Thus, fidelity increases as a spatial representation increases in amplitude, and increases as dispersion – the width of the spatial representation – decreases.

We found that spatial position was represented with high fidelity during the delay period of both memory tasks in several retinotopic ROIs. Representational fidelity tended to peak earlier for the WM task (mean timepoint across ROIs: WM 6.89 seconds after stimulus presentation; LTM 8.89 seconds), likely due to sensory activity evoked by the small dot stimulus marking the to-be-remembered position in the WM task. However, the spatial representation persisted well into the delay period, after the initial sensory response started to decay (**Figure 4C**, V1 – IPS2). In the lateral parietal ROIs (dLatIPS, vLatIPS, AnG), there appeared to be no sensory-evoked response, and no significant evidence of sensory reinstatement either.

In the remainder of our analyses, we report results from data averaged over TRs either early in the delay period (TRs 2, 3, and 4, or 2-6 seconds after stimulus presentation) or late in the delay period (TRs 5, 6, and 7, or 8-12 seconds). **Figure 5A** summarizes representational fidelity late in the delay period (comparable to **Fig. 4C**). As we saw previously, the retinotopic ROIs show evidence of sensory reinstatement, while the lateral parietal cortex (LPC) ROIs do not. To test this, we first ran a two-way repeated measures ANOVA with ROI and memory task as factors and determined statistical significance by comparing the F-scores to a permuted null distribution. We found a main effect of ROI (p < 0.001) and a significant interaction (p = 0.003), but no significant difference between memory tasks (p = 0.191). This interaction suggests that some ROIs have a larger difference between memory tasks than others (e.g., higher WM fidelity in V1 – V3 / V3AB, **Figure 5A**), even though the average difference between tasks is insignificant.

Bayesian t-tests on the data from the late delay period revealed that most ROIs had weak-to-moderate evidence for the hypothesis that there was no difference between WM and LTM (BF_01_ for V3: 1.47, V4: 2.25, IPS0: 1.80, IPS1 2.18; IPS2 2.89; dLatIPS 2.16; vLatIPS 3.24; AnG 2.92), while in the remaining ROIs there was weak evidence for the alternative hypothesis that one condition has higher fidelity (BF_01_ for V1: 0.35, V2: 0.77, V3AB: 0.45; Jeffreys, 1961; Schönbrodt & Wagenmakers, 2018). Overall, we do not find compelling evidence of a difference in sensory reinstatement between model-based representations during WM and LTM.

One possible explanation for this null result in LPC ROIs is that there was poor signal in the sensory mapping task, and this led to unstable estimates of the spatial selectivity of each voxel that did not generalize well to the memory tasks. To evaluate this possibility, we used a leave-one-run-out cross-validation procedure to train and test models based only on data from the sensory mapping task. In this analysis, shown in **Figure 5B**, we found that activation patterns in all retinotopic and non-retinotopic ROIs except vLatIPS contained information about the viewed position of the mapping stimulus (all significant p’s < 0.05). Thus, even the non-retinotopically organized areas dLatIPS and AnG encoded information about the location of a continuously visible stimulus, suggesting that a failure to find information about remembered positions was not due to poor SNR in the mapping task.

In summary, we assessed the fidelity of memory representations using a fixed encoding model that was trained on sensory-evoked responses from an independent task. We found that activation patterns in retinotopic ROIs contained information about the remembered spatial position in both the WM and LTM tasks, and there was little evidence for a difference in representational fidelity between the memory tasks after ∼8 seconds into the delay period. This is a compelling test of the reinstatement hypothesis because it demonstrates fMRI activation patterns evoked by viewing and attending a spatial position have a strong overlap with activation patterns elicited by remembering the same spatial position. In contrast, non-retinotopically organized ROIs in parietal cortex did not contain information about remembered positions in either memory task when the model was trained on the sensory localizer and tested on data from each memory task. Taken together, these data imply that information reinstated from both WM and LTM is encoded in a sensory-like code, but only in retinotopically organized regions of occipital and parietal cortex.

## Model cross-generalization to compare sensory and mnemonic representations in retinotopic and non-retinotopic ROIs

The presence of spatial information in the lateral parietal cortex ROIs during a perceptual task raises the possibility that they do encode spatial information in the memory tasks, but in a *different format* than the sensory evoked code.

To evaluate this possibility, we used a balanced cross-validation procedure to train and test the spatial IEM separately within each memory task (**Figure 5C**). A permuted two-way ANOVA of ROI by task only showed a significant effect of ROI (ROI p < .001; task p = 0.052; interaction p = 0.071). We then tested each ROI and task individually as before (FDR correcting over all comparisons, q = 0.05). All of the retinotopically organized ROIs contained information about the remembered spatial position in both the WM and the LTM tasks (all p’s < 0.005). However, we also found evidence that activation patterns in dLatIPS and AnG also represented the remembered spatial position in both tasks (dLatIPS WM p = 0.002, LTM p = 0.027; AnG WM p < 0.001, LTM p = 0.072). In contrast, vLatIPS did not contain position-specific representations of the remembered position (WM p = 0.165, LTM p = 0.549). The observation of information about remembered position when training and testing was carried out separately for each memory task – but not when we trained on the sensory localizer – suggests that dLatIPS and AnG encode mnemonic information in a format that differs from the sensory evoked response patterns.

Finally, we also performed cross-training between the memory tasks (i.e., train WM, test LTM; train LTM, test WM) to more directly estimate the extent to which memory-related activation patterns were shared across the two tasks (**Figure 5D**). The two-way ANOVA showed a main effect of ROI and a significant interaction (ROI p < .001; task p = 0.211; interaction p < .001). Similar to the interaction effect when training on the independent sensory task, the interaction appeared to be driven by higher fidelity when training on the WM task and testing on the LTM task in the early visual areas (red dots; **Figure 5D**). When we tested each ROI and task individually, we found significant generalization between the WM and LTM tasks in all of the retinotopic ROIs and in dLatIPS and AnG (all p’s < 0.05). Interestingly, we also found significant information in the cross-training analysis in vLatIPS for both memory tasks (WM p < 0.001, LTM p = 0.003). This was surprising given that vLatIPS did not contain information about either task when training was carried out on the sensory mapping task or when training and testing was done entirely within task. We speculate that the significant effect observed when cross-training between the memory tasks was due to the IEM isolating a representation that was general to both memory tasks, and that allowed the model to successfully cross-generalize. In contrast, training on the sensory mapping task or within each memory task may have caused the model to overfit any noise that did not contribute to the mnemonic stimulus representations in vLatIPS.

Overall, these findings suggest that dLatIPS and AnG encode mnemonic information for spatial position during WM and LTM in a different format than the original perceptual representation. Moreover, the WM/LTM cross-training analysis suggests that the representational format of remembered information in dLatIPS, vLatIPS and AnG is at least partially common to both WM and LTM.

### Task classification

In the analyses above we used different IEM training and testing schemes to test whether the format of reinstated memories was shared between WM and LTM. Next, we examined a related but distinct question: is there a discriminable difference between the activation patterns elicited by WM recall and LTM recall? ROIs that share representational codes may still contain encode some information unique to the memory task. To test this possibility, we trained a binary support vector machine (SVM) to classify each memory trial as belonging to the WM or LTM task. The SVM was trained using data balanced across spatial position and task.

We ran the SVM classifier for both the early and late temporal epochs as defined above (early epoch: TRs 2 – 4, late epoch: TRs 5 – 7). We found that we could discriminate between the two tasks early in the delay period across all retinotopic ROIs and in both dLatIPS and vLatIPS (**Figure 6**). We believe this effect is largely driven by the presence of a sensory stimulus in the WM task, but not the LTM task. Note that successful SVM discrimination cannot be driven by differences in univariate delay period activity, because we removed the mean within each memory task before training and testing the classifier (*Methods*). However, we also found that early visual areas (V1 – V4) failed to discriminate between tasks in the late delay period. Instead, only activation patterns in retinotopic parietal regions discriminated between the WM and LTM task during the late temporal epoch. We also found that activation patterns in dorsal and ventral lateral IPS also discriminated between tasks during the late temporal epoch. Lastly, we found that activation patterns in angular gyrus were unable to discriminate between tasks entirely.

**Figure 6.**
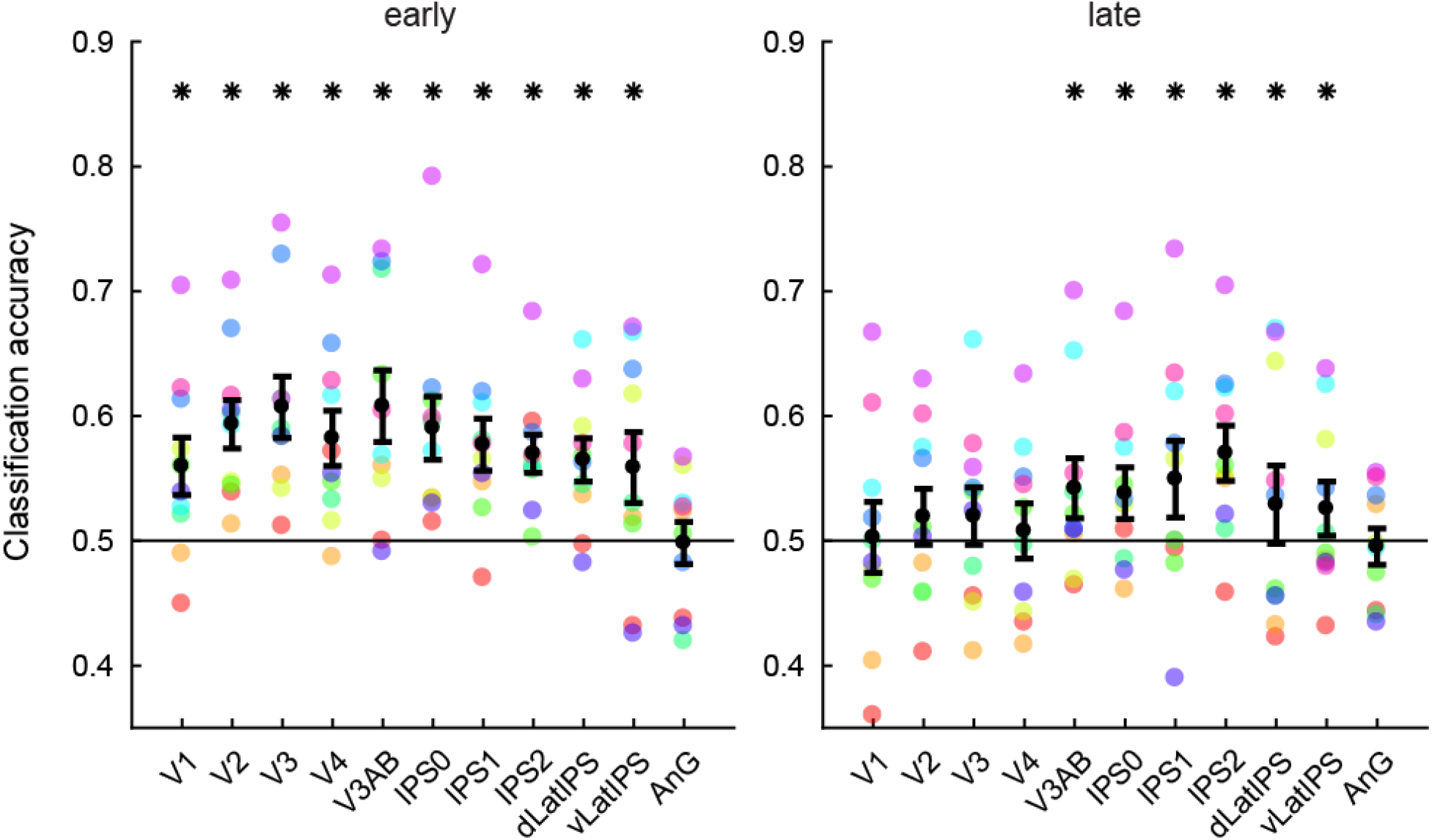
Task decoding in the early and late epochs of the delay period in each ROI. Colored dots show data from each individual subject, and error bars are 95% confidence intervals estimated with bootstrapping. Classifier accuracy is significantly above chance (*) if it passes a one-tailed a permutation test, FDR-corrected across ROI and temporal epoch (q = 0.025). Only activation patterns in areas V3AB and regions of parietal cortex supported above chance decoding accuracy in the late epoch.

To evaluate the differential classification accuracy across ROIs, we ran a two-way repeated measures ANOVA of ROI by temporal epoch on classifier accuracy and calculated p-values as previously described. Both main effects of ROI and epoch were significant (p’s < .001), and there was a significant interaction between the two (p = .001). Since AnG did not show any ability to discriminate between task, we re-ran the ANOVA without this ROI. While there was no main effect of ROI (p = 0.076), the effect of epoch (p < .001) and the interaction (p = 0.01) remained significant. To test the hypothesis that the interaction was driven by the difference in discriminability between epochs, we ran post-hoc one-way ANOVAs of ROI on classifier accuracy separately within each epoch (main effect of ROI on early epoch: p = .065; late epoch: p = 0.016). Since all ROIs were able to discriminate task in the early epoch, we saw no significant effect of ROI there. These data suggest that although the representation of the remembered location is similar between the WM and LTM tasks, in most ROIs that we examined, information that differentiates the two tasks is selectively present in both retinotopic and lateral parietal regions.

## Discussion

The present study, in which we examined WM and LTM using an experimental design that closely matched behavioral precision, makes three main contributions. First, we demonstrate that the fidelity of sensory-like representations is similar between both WM and LTM in both occipital (V1 – V4) and parietal (V3AB, IPS0-2) retinotopic regions. Second, we provide evidence that while lateral parietal regions do not reinstate information in a sensory-like format, they do encode spatial memories in a format that is partially shared between the WM and LTM tasks. Third, we show activity patterns in parietal regions, but not early visual areas, contained information about whether subjects were maintaining information encoded into WM or retrieved from LTM. We propose that these parietal areas may be jointly coding information about the content and the source of visual memories.

The results that directly compare between WM and LTM representations have implications for theories about how memory systems interact with one another. Many theories posit that LTM retrieval places remembered content directly into WM. That is, both WM and actively retrieved LTMs are maintained via the same processes, a capacity-limited state called the “focus of attention” (Atkinson & Shriffin, 1968; Cowan, 1995; D’Esposito & Postle, 2015; Fukuda & Woodman, 2017; LaRocque, Lewis-Peacock, & Postle, 2014). Our finding that WM and LTM representations are both represented in the same retinotopic regions in the same sensory-like format provide compelling evidence for this account.

This theory is also consistent with our finding that lateral parietal regions encode mnemonic information in a format that was shared between WM and LTM tasks. We could robustly reconstruct the remembered position from dLatIPS activity when we trained and tested the model within or across memory tasks. The pattern of results was similar in AnG, although only modest task-specific mnemonic information was observed in the within-task training/testing analysis (see **Table 1**; **Figure 5C**). In contrast, we only observed spatial information in vLatIPS when models were trained on one memory condition and tested on the other. Together, these results support a general role for these regions in representing spatial information in both WM and LTM. However, since the format is not sensory-like, it suggests that the information is somehow reformatted or transformed. This is supported by recent evidence (Favila et al., 2018; Xiao et al., 2017) that have led some to argue that these mnemonic representations are spatially transformed versions of perceptual representations (Favila et al., 2020).

**Table 1.** Summary of effects in a representative set of areas in occipital cortex (V1), retinotopically defined parietal cortex (IPS0) and the lateral parietal ROIs (dLatIPS, vLatIPS, AnG). A “1” indicates a significant effect and a “0” indicates a non-significant effect.

| Method | Test of representation | V1 | IPS0 | dLatIPS | vLatIPS | AnG |
| --- | --- | --- | --- | --- | --- | --- |
| IEM | Sensory info (control) | 1 | 1 | 1 | 0 | 1 |
| IEM | Sensory-like reinstatement | 1 | 1 | 0 | 0 | 0 |
| IEM | Task-specific | 1 | 1 | 1 | 0 | ~1 |
| IEM | Task-general (cross-training) | 1 | 1 | 1 | 1 | 1 |
| SVM | Task discriminable | 0 | 1 | 1 | 1 | 0 |

This transformation may be functional, ensuring that information is memory is distinguishable from incoming sensory information (Bettencourt & Xu, 2015; Stokes, 2015; Xu, 2017). We note this is purely speculative in the absence of additional experimental data evaluating memory fidelity in the face of different kinds of competing distractors (i.e. does the degree to which the codes in lateral parietal cortex are transformed away from sensory codes predict success in the face of concurrent sensory distractors?).

We also found that activity patterns in almost all parietal ROIs, but not occipital ROIs, could decode whether a trial belonged to the WM or LTM task (**Figure 6**). The one exception was AnG, which could not discriminate between the two memory tasks. We discuss three possible reasons we observed this finding in the remaining parietal ROIs. One possibility is that parietal regions encode the task set (i.e., higher-level representations that govern the execution of a specific task). Indeed, past work has shown that parietal regions encode higher-order task representations, such as the task rule that the subject is executing (Bode and Dylan-Haynes, 2009; Woolgar et al., 2011a; 2011b). The second possibility is that the WM and LTM tasks require different attentional processes. Parietal regions have been reported to be engaged during attention tasks, WM tasks, and LTM tasks (Awh & Jonides, 2001; Awh, Vogel, & Oh, 2006; Cabeza, Ciaramelli, Olson, & Moscovitch, 2008; Hutchinson et al., 2014; Sestieri, Shulman, & Corbetta, 2017). Researchers have separately proposed that the parietal mechanisms involved in spatial attention also support spatial WM (Awh & Jonides, 2001; Awh et al., 2006), and that parietal regions mediate attention to long-term memories during retrieval (Cabeza et al., 2008; Sestieri et al., 2017). The ability of the SVM decoder to distinguish between WM and LTM tasks in parietal regions raises the possibility that these attention-mediated memory processes may not be identical across tasks. The last possibility is that parietal regions encode the *source* of memory representations (perception for WM, and an internal store for LTM). One issue with relying solely on cortical reinstatement for memory recall is that multiple sensory representations in one cortical region, such as area V1, could be attributable to ongoing perception, recent perception (WM), or distant perception (LTM). In order to maintain separable representations within sensory areas, parietal regions may additionally represent the source of sensory information. This could help protect reinstated sensory memories from interference by concurrent sensory input (Bettencourt & Xu, 2015; Rademaker et al., 2019). Further work is needed to test these hypotheses.

Finally, we note that our investigation of memory representations is limited to visual and parietal cortex, and does not provide any evidence about the differential role of other areas, such as the medial temporal lobe (MTL), which is thought to play a role in both LTM and WM (Squire & Wixted, 2011). Although the MTL has long been linked with LTM (Mishkin, 1978; Scoville & Milner, 1957; Squire & Zola-Morgan, 1978) a review of existing evidence suggests that the hippocampus is necessary for some types of WM tasks – perhaps specifically when they are complex (Yonelinas, 2013) or spatial tasks (Nadel & Hardt, 2011). Future studies that are optimized for imaging and segmenting the MTL could use similar techniques to compare memory reinstatement and retrieval across WM and LTM in MTL regions.

In conclusion, our findings suggest that remembered spatial position encoded into WM or retrieved from LTM are represented similarly in occipital and parietal cortex. In retinotopic regions, activity patterns recapitulated those that were observed during an independent sensory stimulation condition, showing that a sensory recruitment model provides a useful perspective for how visual details are maintained, regardless of the specific memory system. In non-retinotopic areas of lateral parietal cortex, mnemonic representations appear to share a sensory format across WM and LTM that is distinct from a sensory-like code. Moreover, parietal regions – both retinotopic and lateral – also represented the memory task being performed, unlike retinotopic occipital regions.

## Acknowledgements

Funding provided by NIH-MH087214 to EA and NIH-EY025872 to JTS, NSF GRFP to VAV, and F32-EY028438 and Sloan Research Fellowship to TCS. We thank Megan deBettencourt and Nuttida Rungratsameetaweemana for helpful discussions.

